# *oca2* targeting using CRISPR/Cas9 in the Malawi cichlid *Astatotilapia calliptera*

**DOI:** 10.1101/2022.01.01.474687

**Authors:** Bethan Clark, Joel Elkin, Aleksandra Marconi, George F. Turner, Alan M. Smith, Domino Joyce, Eric A. Miska, Scott A. Juntti, M. Emília Santos

## Abstract

Identifying genetic loci underlying trait variation provides insights into the mechanisms of diversification, but demonstrating causality and characterising the role of genetic loci requires testing candidate gene function, often in non-model species. Here we establish CRISPR/Cas9 editing in *Astatotilapia calliptera*, a generalist cichlid of the remarkably diverse Lake Malawi radiation. By targeting the gene *oca2* required for melanin synthesis in other vertebrate species, we show efficient editing and germline transmission. Gene edits include indels in the coding region, likely a result of non-homologous end joining, and a large deletion in the 3′ UTR due to homology-directed repair. We find that *oca2* knock-out *A. calliptera* lack melanin, which may be useful for developmental imaging in embryos and studying colour pattern formation in adults. As *A. calliptera* resembles the presumed generalist ancestor of the Lake Malawi cichlids radiation, establishing genome editing in this species will facilitate investigating speciation, adaptation and trait diversification in this textbook radiation.

## Introduction

Identifying the genetic and developmental mechanisms underlying novel and variable morphologies is key to understanding how diversity arises in nature. Instances of adaptive radiation, that is, the rapid formation of an abundance of diverse species from a common ancestor, are perfect systems to delve into the basis of diversification and adaptation to distinct ecological niches (Schluter, 2000). Cichlid fishes are a textbook example for such adaptive radiations. They are one of the most species-rich vertebrate families comprising over 2200 species which exhibit extraordinary morphological, physiological and behavioural variation (Kocher, 2004; Salzburger, 2018; Santos and Salzburger, 2012). The majority of species (~2000) are found in the East African lakes, Tanganyika, Victoria and Malawi. Lake Malawi alone has over 800 species that emerged in the last 800 000 years (Malinsky et al., 2018; Salzburger, 2018). They show extensive morphological variation in body shape, craniofacial skeleton, jaw apparatus, lateral line system, brain, vision and pigmentation phenotypes among other traits (Carleton et al., 2016; Edgley and Genner, 2019; Kratochwil et al., 2018; Powder and Albertson, 2016; Roberts et al., 2017; Ronco et al., 2021; Santos et al., 2014; Sylvester et al., 2010). Despite their morphological diversity, the average sequence divergence between Malawi cichlid species pairs is only 0.1-0.25%, thus within this lake the evolution of divergent phenotypes seems to occur through comparatively minor genetic changes (Malinsky et al., 2018; Svardal et al., 2020). Their genetic similarity enables interspecific hybridisation, which can be used for quantitative trait loci (QTL) analysis to uncover genes underlying variation in species specific traits. This is bolstered by their amenability to the lab and the wealth of genomic resources that have been made available in recent years, including many representative reference genomes (Brawand et al., 2014). While the aforementioned tools facilitate the discovery of loci associated with trait diversification, proof of causality can only be achieved by testing candidate gene function through genome editing.

Here, we report the application of CRISPR/Cas9 to generate coding and non-coding sequence mutants in the cichlid *Astatotilapia calliptera*, a maternal mouthbrooder cichlid fish that is part of the Malawi haplochromine radiation. *A. calliptera* occupies a rich diversity of habitats, including Lake Malawi, as well as peripheral rivers and lakes (Parsons et al., 2017). Phylogenetic analysis shows that all Malawi cichlid species can be grouped into seven eco-morphological groups, resulting from three separate cichlid radiations that stemmed from a generalist *Astatotilapia-type* ancestral lineage (Malinsky et al., 2018). As such, *A. calliptera* is a useful model in which to develop functional tools to explore Malawi cichlid speciation and adaptation. We specifically focused on one *A. calliptera* population from a small crater lake situated north of Lake Malawi (Figure 1A) referred to in the literature as Lake Masoko (variant spelling Massoko, as used by the German colonial administration) and known locally as Lake Kisiba (Turner et al., 2019). *A. calliptera* from Lake Masoko/Kisiba is at an early stage of adaptive divergence where two diverging ecomorphs differ in body shape, diet, trophic morphology and body colouration (Figure 1B and 1C) making it also an ideal system to study the early stages of speciation. Importantly, *A. calliptera* has a high-quality reference genome and is amenable to the lab environment. They have a 8-12 month generation time, breeding readily in a non-seasonal fashion allowing for year-round egg collection for gene editing and embryonic developmental studies.

**Figure 1:**
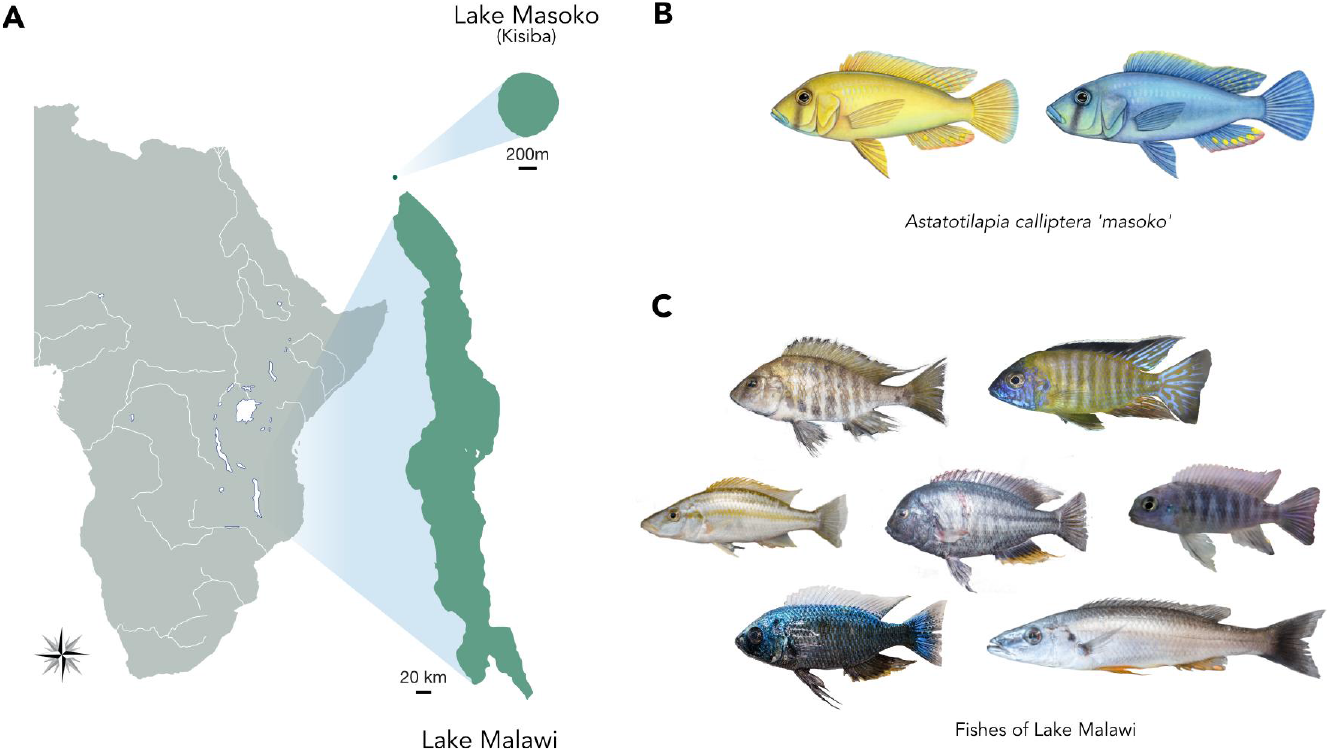
*Astatotilapia calliptera* and Malawi cichlids. **A)** Geographical location of Lake Malawi and Lake Masoko/Kisiba. Map diagram courtesy of Gregóire Vernaz. **B)** The two *A. calliptera* eco-morphs from Lake Masoko/Kisiba. Drawings by Julie Jonhson. **C)** A snapshot of the diversity of forms present in Lake Malawi. Photographs courtesy of Hannes Svardal.

We chose to generate mutants for the *Oculocutaneous albinism type II* gene (*oca2*). *Oca2* is a relatively well-characterised gene and has a readily visible phenotype where black pigment production - melanin - is impaired (Beirl et al., 2014; Klaassen et al., 2018). It encodes a melanosomal transmembrane protein associated with the intracellular trafficking of tyrosinase, a rate-limiting enzyme in the melanin biosynthesis pathway. *Oca2* has been associated with the evolution of amelanism and albinism in natural populations in multiple vertebrate species, such as humans, snakes, cavefish and cichlids (Edwards et al., 2010; Kratochwil et al., 2019; O’Gorman et al., 2021; Protas et al., 2006; Saenko et al., 2015).

In ray-finned fishes, pigmentation patterns are generated by the different number, combinations and arrangement of pigment cells: such as black melanophores, yellow to red xanthophores and reflecting silvery iridophores (Parichy, 2021). All pigment cell classes share an embryonic origin, deriving from the neural crest cell population during early development. Pigmentation patterning has been extensively studied in zebrafish, where the adult pigment pattern emerges through the migration and interaction between pigment cells, as well as interactions between the cells and the tissue environment (Kelsh and Barsh, 2011; Parichy and Spiewak, 2015; Singh and Nüsslein-Volhard, 2015). *Oca2* knockout in zebrafish is known to impair melanin production, melanophore differentiation and survival, as well as increasing the abundance of iridophores without affecting adult patterning (Beirl et al., 2014). Importantly, *oca2* mutants are viable, making *oca2* single guide RNA (sgRNA) micro-injections an ideal tool to assess rates of mutagenesis and germline transmission, and to establish CRISPR/Cas9 protocols in *A. calliptera*.

CRISPR/Cas9 editing tools have revolutionised gene function analysis in a multitude of non-model species. This is due to the simplicity of the system which requires only Watson-Crick base pairing between a sgRNA and its target sequence. The Cas9 protein will form a complex with the sgRNA, which will recognise and bind to its target sequence. The Cas9 protein will then induce a double-stranded DNA break. When double-stranded breaks are formed, the intrinsic cellular repair machinery will put the two back together either using non-homologous end joining (NHEJ) or homology-directed repair (HDR). NHEJ is an imprecise mechanism that generates small insertions or deletions that result in a frameshift. This method has been widely used to induce frameshift mutations in coding protein sequences leading to loss of function alleles in many model systems including other cichlids (Fang et al., 2018; Höch et al., 2021; Kratochwil et al., 2018; Li et al., 2021, 2014; Livraghi et al., 2021; Rasys et al., 2019; Wang et al., 2021). HDR on the other hand utilises a DNA template to guide the repair, thus by providing a single-stranded DNA (ssDNA) template, one can either insert a sequence of interest (e.g. allelic exchange) or generate larger and more precise deletions (Hisano et al., 2015; Kimura et al., 2014; Li et al., 2019; Wierson et al., 2020).

Here, CRISPR/Cas9-mediated knockout was employed using these two approaches - NHEJ and HDR - to respectively target coding and non-coding regions. First, exons 1 and 3 of the *oca2* coding sequence were targeted with sgRNAs with the intent of generating frameshift coding mutations. Second, an HDR ssRNA template was used to generate a ~1,100 bp deletion in the 3′ untranslated region (UTR) of *oca2*. More specifically, two sgRNA target sequences were identified – one at either end of a ~1,100 bp region in the 3′ UTR. These were then co-injected with a 100 bp ssDNA DNA template that was mutually homologous to 50bp at either target site. This ssDNA template facilitates homology directed repair (HDR) to replace the region in between the target sites with a deletion (Li et al., 2019).

Using these two approaches, we generated coding and non-coding *oca2 A. calliptera* mutants using site directed disruption with CRISPR/Cas9. As expected, loss of *oca2* function results in amelanism due to the inability to synthesise melanin. The deletion in the 3′ UTR region yielded no visible phenotypic effect. Coding and non-coding mutations were successfully transmitted to the next generation. The establishment of CRISPR/Cas9 methodologies in *A. calliptera* provides a platform for the future analysis of coding and regulatory variation in one of the most astonishing vertebrate adaptive radiations - Malawi cichlid fishes - and will enhance our understanding of the genomic basis of organismal diversity.

## Results

### Site-directed disruption of *A. calliptera* oca2 coding sequence

To demonstrate the feasibility of genome editing in *A. calliptera*, we generated mutations in the coding sequence of *oca2*. Two injection mixes were used, each containing two sgRNAs with the intent to increase the chances of introducing mutation (Li et al., 2021). These were co-injected with Cas9 protein into fertilised single-cell eggs (Figure 2A, Table S1). The first mix contained sgRNA 1 and sgRNA 2 respectively targeting exon 1 and exon 3 (Figure 2A). The second contained sgRNA 3 and sgRNA 4 both targeting exon 1 (Figure 2A). Exons near the 5′ end of the gene were selected to increase the chance of a frameshift mutation causing a missense translation for most of the length of the protein sequence. Embryos were screened at 4 days post fertilisation (dpf, 25C) when the retinal pigment epithelium becomes pigmented with melanin. Both injection mixes yielded mosaic individuals with an average survival rate of 28.5%. The percentage of mosaic individuals was variable ranging from 18% to 100% with an average of 54% (Table 1).

**Figure 2:**
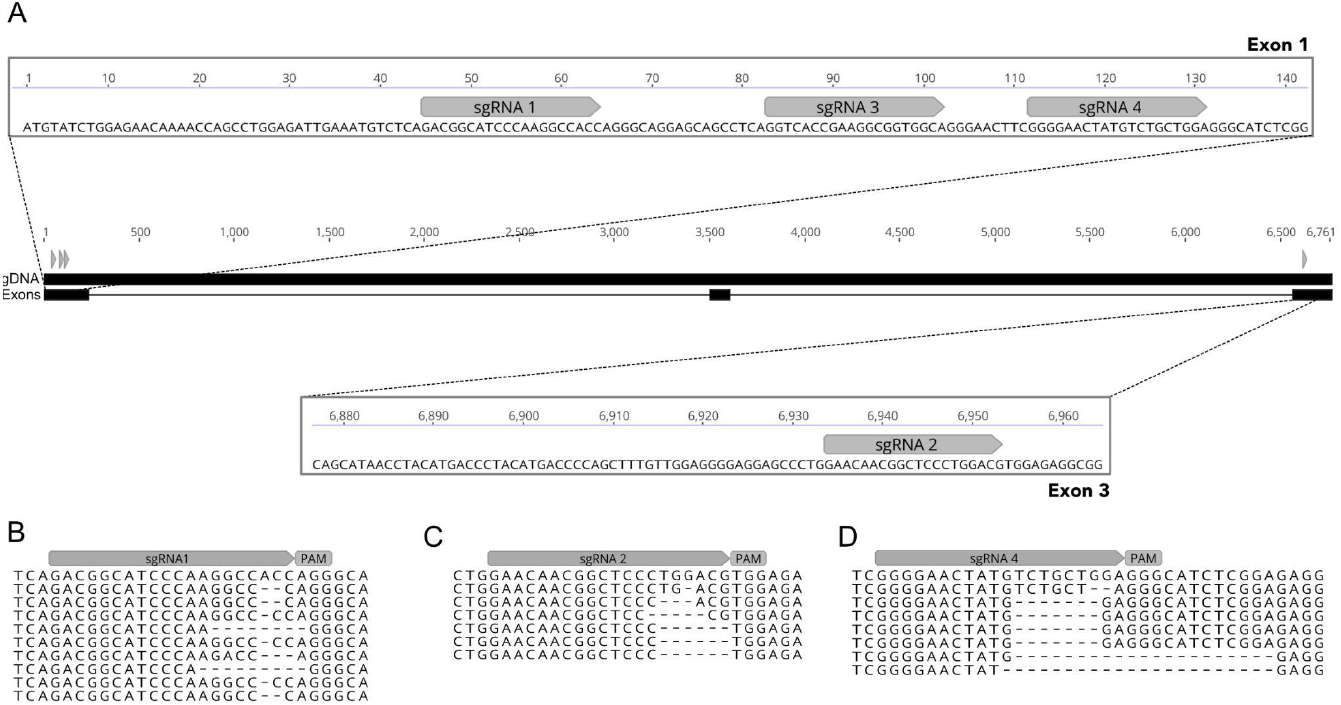
Efficient indel generation of *oca2* coding sequence by Crispr/Cas9. **A)** Four sgRNAs were designed to cut the genomic sequence at exon 1 and exon 3. Two injection mixes were used, one containing sgRNA1 and sgRNA2 targeting exon 1 and exon 3, and the other containing sgRNA3 + sgRNA 4 both targeting exon 1. Alignment of mutant F1 individuals derived from crosses #1, #3 and #4 are shown for **B)** sgRNA1, **C)** sgRNA2 and **D)** sgRNA4. All F1 individuals were wild-type at the cut site of sgRNA3.

**Table 1:**
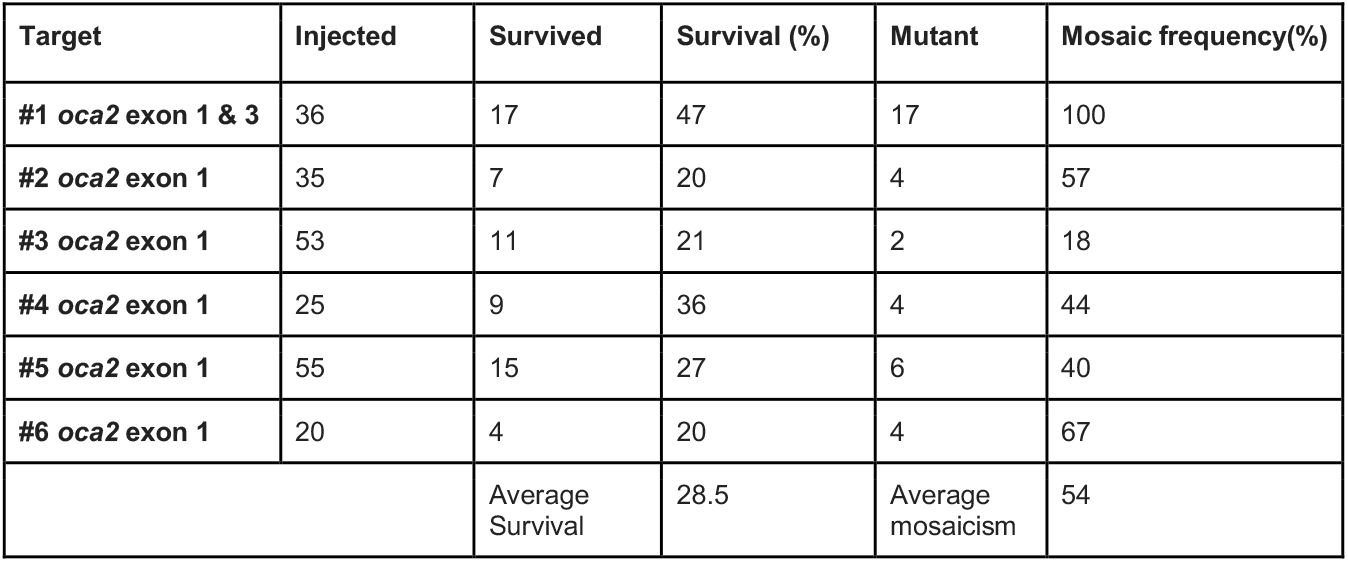
Percentage of *oca2* mosaic individuals induced by CRISPR/Cas9 in G0s.

| Target | Injected | Survived | Survival (%) | Mutant | Mosaic frequency(%) |
| --- | --- | --- | --- | --- | --- |
| #1 *oca2* exon 1 & 3 | 36 | 17 | 47 | 17 | 100 |
| #2 *oca2* exon 1 | 35 | 7 | 20 | 4 | 57 |
| #3 *oca2* exon 1 | 53 | 11 | 21 | 2 | 18 |
| #4 *oca2* exon 1 | 25 | 9 | 36 | 4 | 44 |
| #5 *oca2* exon 1 | 55 | 15 | 27 | 6 | 40 |
| #6 *oca2* exon 1 | 20 | 4 | 20 | 4 | 67 |
|  |  | Average Survival | 28.5 | Average mosaicism | 54 |

To investigate if mutations are transmitted to the following generations, fish showing mosaic phenotypes were raised to adulthood. From these G0 adult fish, 4 males were crossed with WT females. In addition, we incrossed one *oca2* mosaic male with two *oca2* mosaic females to obtain *oca2* mutants carrying two *oca2* coding knock-out alleles and hence with a visible amelanistic phenotype in one generation (Table 2). The genotyping of sequencing products derived from the progeny of crosses of male founder individuals with wild-type females showed an average transmission rate of 49% (Table 2). The lowest transmission rate (none) was detected in a male with low levels of phenotypic amelanistic mosaicism, whereas transmission was highly effective for the other three mosaic males with extensive amelanism (40-78%). As germline transmission was high, we were able to generate two incrosses between *oca2* mosaic mutants that both generated amelanistic phenotypes. One of the incrosses generated 10 embryos all with amelanistic phenotypes. Since this mutation is recessive, this result indicates that both progenitors exhibited a transmission rate of 100% for this clutch, generating progeny carrying two *oca2* coding knock-out alleles. The second incross generated 10 embryos with only 2 showing the amelanistic phenotype, which implies a rate of 20% transmission for one of the progenitors. Taken together, germline transmission was high and observed in all crosses where founders showed high levels of phenotypic amelanistic mosaicism (Table 2).

**Table 2:** Germline transmission of 6 founder crosses for the oca2 loss of function mutations.

| Group | Founder cross | #F1s tested | Positive individuals | Germline transmission (%) |
| --- | --- | --- | --- | --- |
| ***oca2* mosaic male x with WT females: Germline transmission quantified by PCR genotyping** |  |  |  |  |
| #1 *oca2* exon 1 | *oca2*<sup>♂1</sup> x wt<sup>♀</sup> | 9 | 7 | 78 |
| #2 *oca2* exon 1 & 3 | *oca2*<sup>♂2</sup> x wt<sup>♀</sup> | 9 | 0 | 0 |
| #3 *oca2* exon 1 & 3 | *oca2*<sup>♂3</sup> x wt<sup>♀</sup> | 9 | 7 | 78 |
| #4 *oca2* exon 1 & 3 | *oca2*<sup>♂4</sup> x wt<sup>♀</sup> | 9 | 3 | 33 |
|  |  |  | Average transmission | 28 |
| **Mosaic *oca2* x Mosaic *oca2*: Germline transmission quantified by presence of amelanistic individuals** |  |  |  |  |
| #5 *oca2* exon 1 | *oca2*<sup>♂5</sup> x *oca2*<sup>♀1</sup> | 10 | 10 | 100 |
| #6 *oca2* exon 1 | *oca2*<sup>♂5</sup> x *oca2*<sup>♀2</sup> | 10 | 2 | 20 |
|  |  |  | Average transmission | 60 |

Sequence analysis of the F1 progeny resulting from crosses #1, #3 and #4 (Table 2) shows that both injection mixes resulted in deletions of variable sizes ranging from 1 bp to 21 bp deletions (Figure 2B, C and D). The F1 progeny from cross #3 and #4, resulting from microinjections using sgRNA1 and sgRNA2, that respectively target exon 1 and exon 3 (Figure 2B and C), show that these two guides have different germline transmission rates, with sgRNA1 presenting a higher frequency (67% for #3 & 33% #4) than sgRNA2 (44% for #3 & 22% for #4) (Supplementary File 1). For cross #3 the transmission rates for each individual guide are lower than the calculated rate for when the two are combined (Table 1, cross #3 and #4), showcasing the benefits of injecting several guides in combination. This is further strengthened by the genotyping results from cross #1, which shows that only one of the two guides injected resulted in indels (sgRNA 4, Figure 2D).

### Deletion of *A. calliptera oca2* 3′ untranslated region

The majority of a given organismal genome is non-coding in nature and regulates the timing and location of gene expression and transcript stability. It has been repeatedly shown that non-coding sequence divergence contributes greatly to cichlid diversity (Baldo et al., 2011; Brawand et al., 2014; Kratochwil et al., 2018; O’Quin et al., 2011). For example, the comparison of the first five cichlid reference genomes showed an abundance of non-coding element divergence and found that transposable element insertions upstream of transcription start sites were associated with expression divergence (Brawand et al., 2014). Further, 3′ UTRs also act as key regulators of gene expression, containing binding sites for microRNAs and RNA-binding proteins (Mayr, 2017). The investigation of cichlid microRNA genes detected signatures of divergent natural selection in microRNA target sites among Lake Malawi cichlids (Loh et al., 2011). A comparative transcriptome analysis has further revealed little divergence at protein-coding sequences, but a high diversity in UTRs (Baldo et al., 2011). Taken together, these studies suggest that regulatory evolution plays a key role in cichlid diversification. Thus, it is important to establish a protocol that allows for testing of the function of non-coding regions associated with trait variation. For this purpose, we took advantage of the HDR CRISPR/Cas9 method to generate a large deletion in the 3′ UTR region of *oca2*.

First, two sgRNA target sequences were identified - one at either end of a 1,096 bp region (Figure 3A). Then, a 100 bp ssDNA repair template was designed to be homologous to the flanking regions of each target site (50 bp upstream and 50 bp downstream), in order to mimic a deletion when compared to the wild-type sequence (Figure 3A). A mix of the two sgRNAs, the ssDNA together with Cas9 protein was microinjected into fertilised single-cell eggs (Table S1). As there was no observable phenotype, the number of mosaic 3′ UTR deletion mutants was assessed by PCR and Sanger sequencing using two primers flanking the cut sites (207 bp upstream the left cut site and 400bp downstream of the right cut site) (Table S2). This assay differentiates between the wild-type individuals and individuals carrying the desired deletion. PCR on wild-type individuals results in only one PCR fragment, whereas mosaic individuals carrying the deletion will show two fragments - the wild-type sequence (~1,715 bp) and the sequence containing the deletion (~624 bp). Using this assay, we determined that the percentage of mosaic individuals was 25% and 0% in the two clutches injected, with the presence of only one positive mosaic mutant (Table 3).

**Figure 3:**
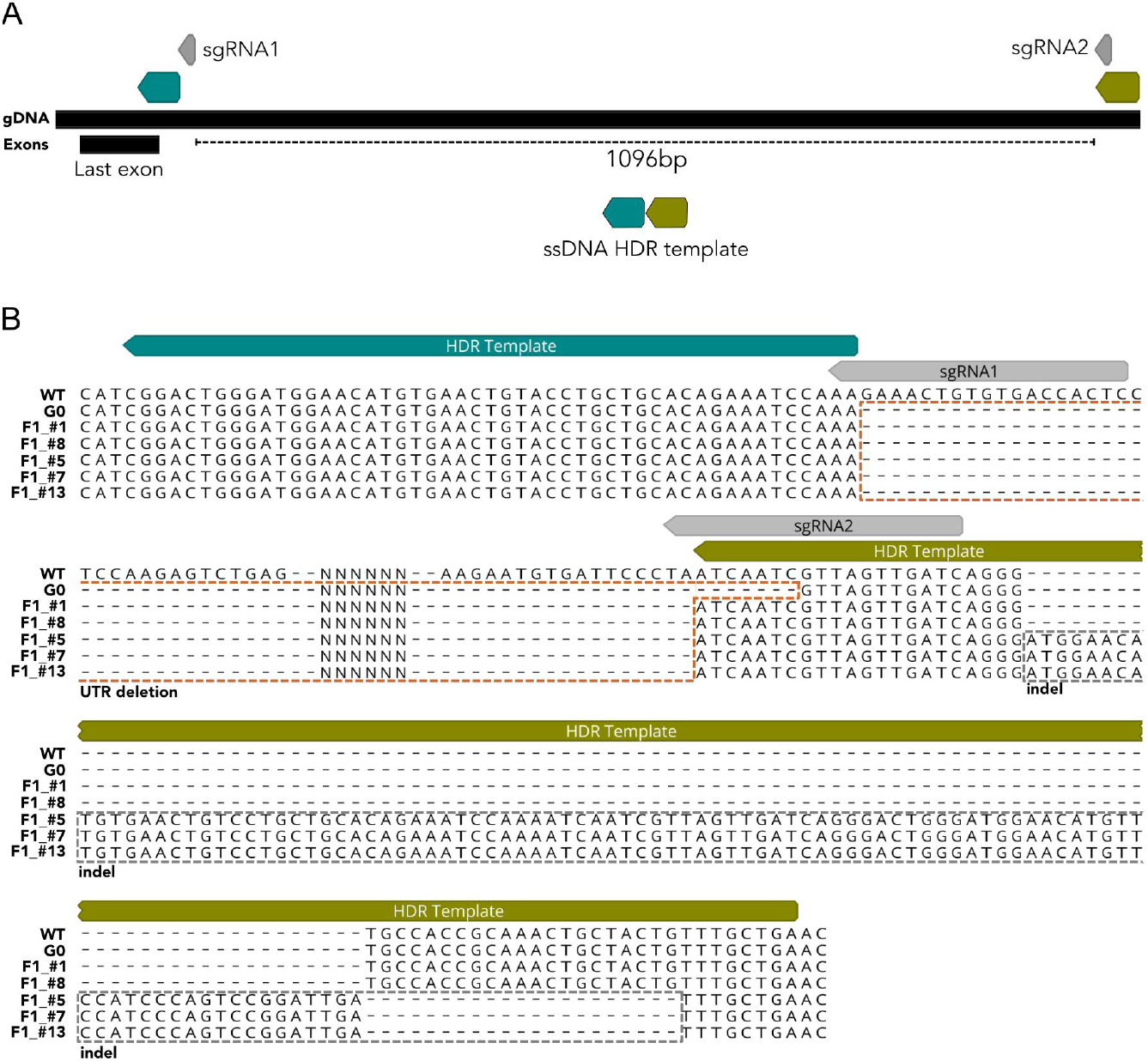
Large deletion of *oca2* 3′ UTR sequence (~1096bp) using one pair of sgRNAs (grey boxes) and one ssDNA HDR template. **A)** A single stranded HDR template with 50 bp left and right homology arms (blue and green boxes respectively) were co-injected with two sgRNAs flanking the desired deletion sites. **B)** Sequencing of the PCR products confirmed the deletion in the G0 and F1s. The 3′ UTR deletion is marked with an orange dashed box. In three of the F1s (#5, #7 and #13) the deletion is followed by a downstream indel marked with a grey dashed box.

**Table 3:** Percentage of *oca2* 3′ UTR mosaic mutants induced by CRISPR/Cas9 in G0s.

| Target | Injected | Survived | Survival (%) | Mutant | Mosaic frequency(%) |
| --- | --- | --- | --- | --- | --- |
| #1 *oca2* 3' UTR | 20 | 4 | 20 | 1 | 25 |
| #2 *oca2* 3' UTR | 58 | 3 | 10 | 0 | 0 |
|  |  | Average Survival | 15 | Average mosaicism | 12.5 |

To determine if deletions are transmitted to the following generations, the G0 mosaic individual for the deletion was raised to adulthood. This *oca2* 3′ UTR mosaic mutant male was crossed with wild-type individuals and showed a germline transmission of 38% (Table 4). We genotyped F1s deriving this cross and confirmed the presence of germline transmission for the deletion. Sequencing of G0 and F1 individuals further confirmed the presence of deletions between the two target sites. The G0 founder shows the presence of a deletion and 5 out of 13 F1s inherited mutations (Figure 3B). While two of the F1s (F1_#1 and F1_#8) have a precise deletion, the other three show the deletion (F1_#5, F1_#7 and F1_#13) followed by a ~100bp downstream insertion (Figure 3B). This insertion shows homology to the HDR template which was potentially inserted - knocked-in - as part of the repair mechanism (Figure S1). These results show that the deletion of large non-coding fragments was successful in *A. calliptera*, but careful screening and sequencing of F1s is required to confirm the presence of precise nature of the deletions. The injection of different sgRNA and HDR template combinations, using larger clutches and screening more F1s will contribute to the refinement of this technique.

**Table 4:**
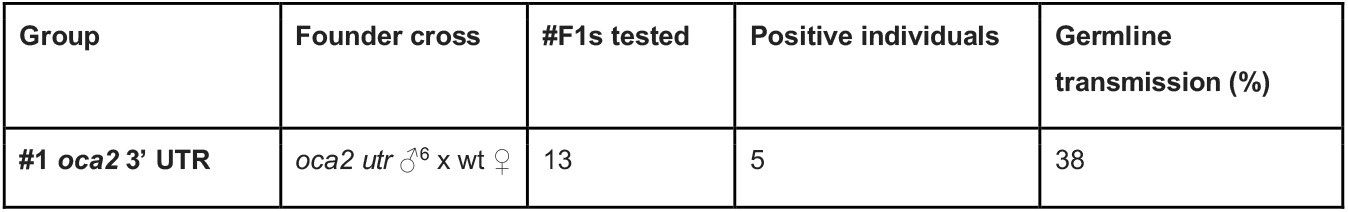
Germline transmission of one founder cross for the *oca2* 3′ UTR deletion.

### Phenotype of *oca2* coding and non-coding mutants in *A. calliptera*

In agreement with previous work in other model systems, *oca2* loss of function mutations led to a reduction in melanic pigmentation. In wild-type embryos, yolk melanophores are the first visible pigment cells to appear on the embryo (Figure 4A), and they remain on the yolk until body wall closure. As such, the first observed phenotype in embryos injected with sgRNAs targeting the *oca2* coding sequence was a reduction in visible yolk melanophore abundance at 4 days post fertilisation (dpf) (Figure 4B). In F1 fish with two *oca2* mutant alleles, there is a complete lack of pigmented melanophores at this stage (Figure 4C); this amelanic phenotype persists throughout development.

**Figure 4:**
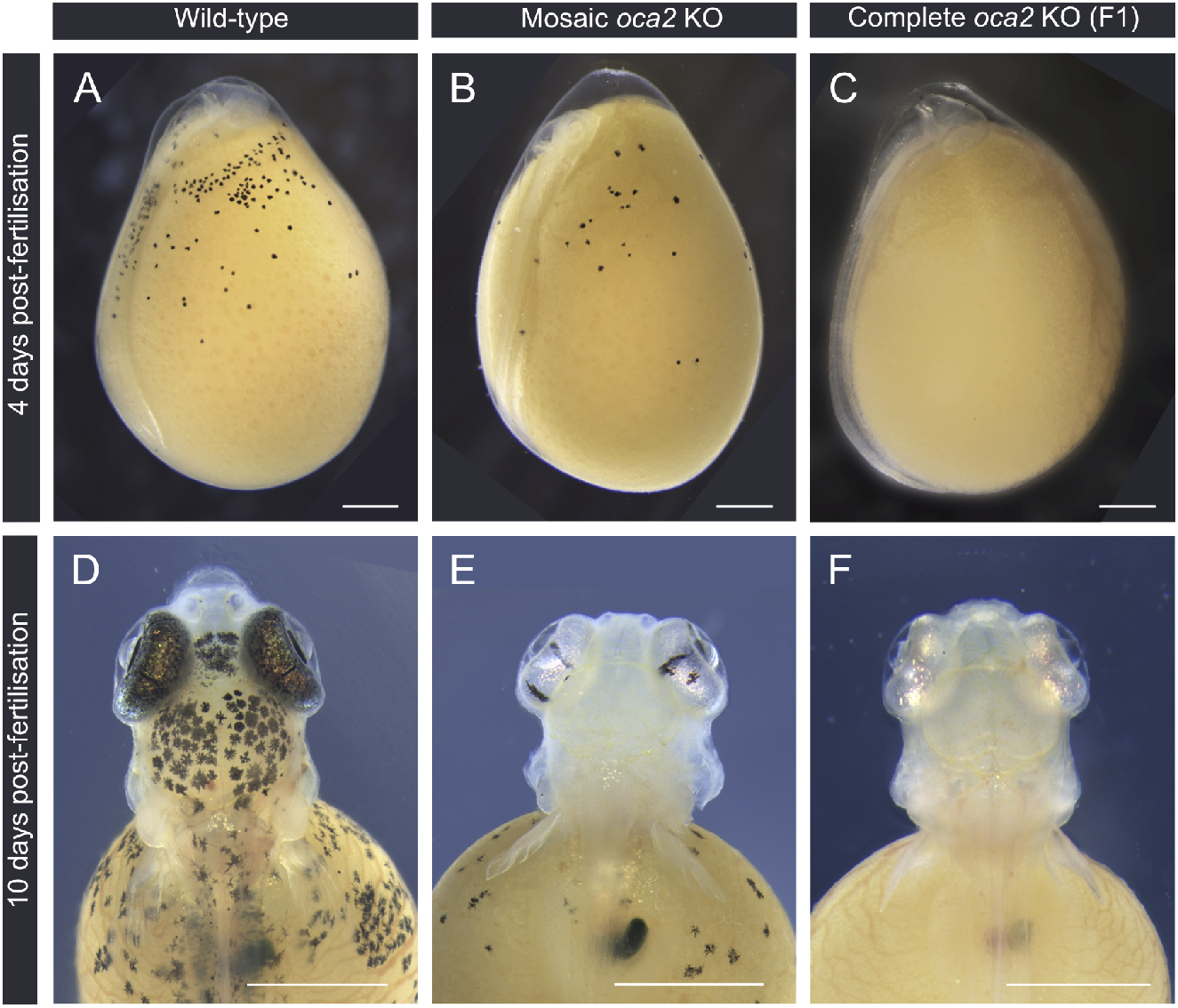
*oca2* loss of function embryonic phenotypes (25° C). **A-C)** Embryo development at 4 days post fertilisation and; **D-F)** 10 days post fertilisation for wild-type (A, D), mosaic G0 (B, E), and F1 *oca2* coding knock-out embryos (C, F). Scale bar 1 mm.

By 10 dpf the degree of melanic coverage along the body increases in wild-type larvae, particularly in the head region, where the yolk melanophores increase in number and are more densely packed. The retina becomes fully pigmented, harbouring both melanophores and iridophores (Figure 4D). On the contrary, *oca2* mosaic mutant larvae continue to have fewer visible melanophores appearing on either the head or on the yolk (Figure 4E) and none in F1 fish with two mutant alleles (Figure 4F). Despite the lack of melanin pigment, the retinae of *oca2* mutant larvae are bright and iridescent indicating the presence of iridophores (Figure 4E and 4F).

Throughout all stages of development described, there is no apparent difference in phenotype between wild-type embryos and the *oca2* 3′ UTR deletion mosaic (Figure S2). Mosaic *oca2* coding knock-outs mutants continue to display a hypopigmented phenotype as adults (Figure 5) while the mosaic *oca2* 3′ UTR deletion mutant has a wild-type phenotype (Figure S3). This result has to be taken with caution as only one G0 individual tested positive for the deletion (Table 3) and may carry very few cells with biallelic mutations. The generation of homozygous mutants is required to fully comprehend the phenotypic effects of the 3′ UTR deletion.

**Figure 5:**
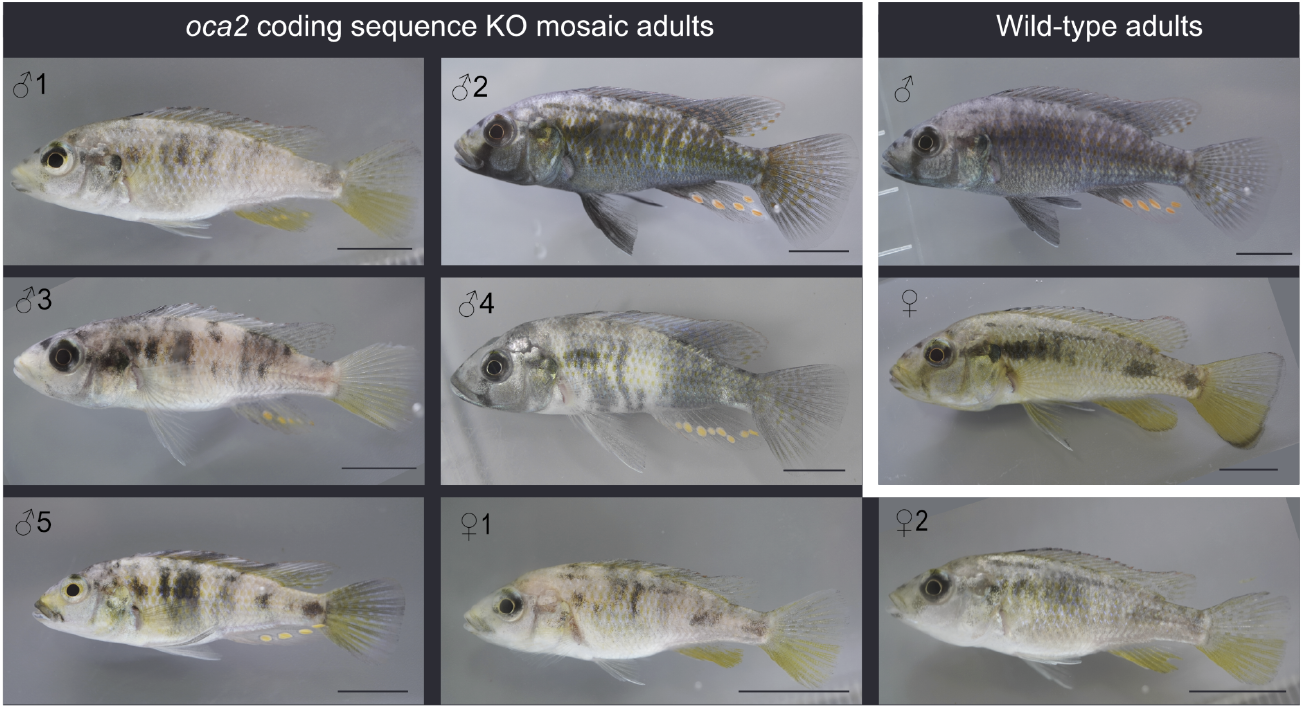
*oca2* coding sequence G0 adult phenotypes. Mosaic injected *oca2* coding region knock-outs compared to wild-type adults. Individual numbering corresponds to Table 2. Scale bar 1 cm

F1 adult fish with two *oca2* coding knock-out alleles have a typical amelanistic phenotype, with a complete absence of black pigmentation (Figure 6). We refer to this phenotype as amelanistic rather than albinistic because albino individuals lack both melanin and other pigments, though this is inconsistently applied across vertebrate taxa (Kratochwil et al., 2019). We compared the pigmentation patterns to the siblings of amelanistics: as the offspring of a cross between two mosaic mutant fish, these siblings may either be homozygous wild-type or heterozygous for the *oca2* coding knock-out which also has a wild-type phenotype (Beirl et al., 2014). In amelanistic adults, lack of pigmented melanophores in the eyes gives the retina a red appearance and black pigmentation patterns on the body and fins are absent, which includes the faint vertical bars on the trunk (Figure 6 A-B), solid black patch on the operculum (Figure 6 C-D), the black bar across the eye in males (Figure 6 C-D), the skin throughout the trunk (Figure 6 E-F), the anterior of the dorsal fin (Figure 6 G-H), and the base, spines, and edges of the caudal fin (Figure 6 I-J). Across the whole body there is greater yellow/orange pigmentation visible and exclusively black areas in wild-type are instead bright and reflective in the amelanistic individuals. The close-up pattern on the body of alternating light and dark patches is maintained in amelanistic adults (Figure 6 E-F). Interestingly, in this pattern the blue regions of the wild-type appear white in the amelanistic, and red erythrophores present in the amelanistic individuals are not visible in the wild-type (Figure 6 E-F).

**Figure 6:**
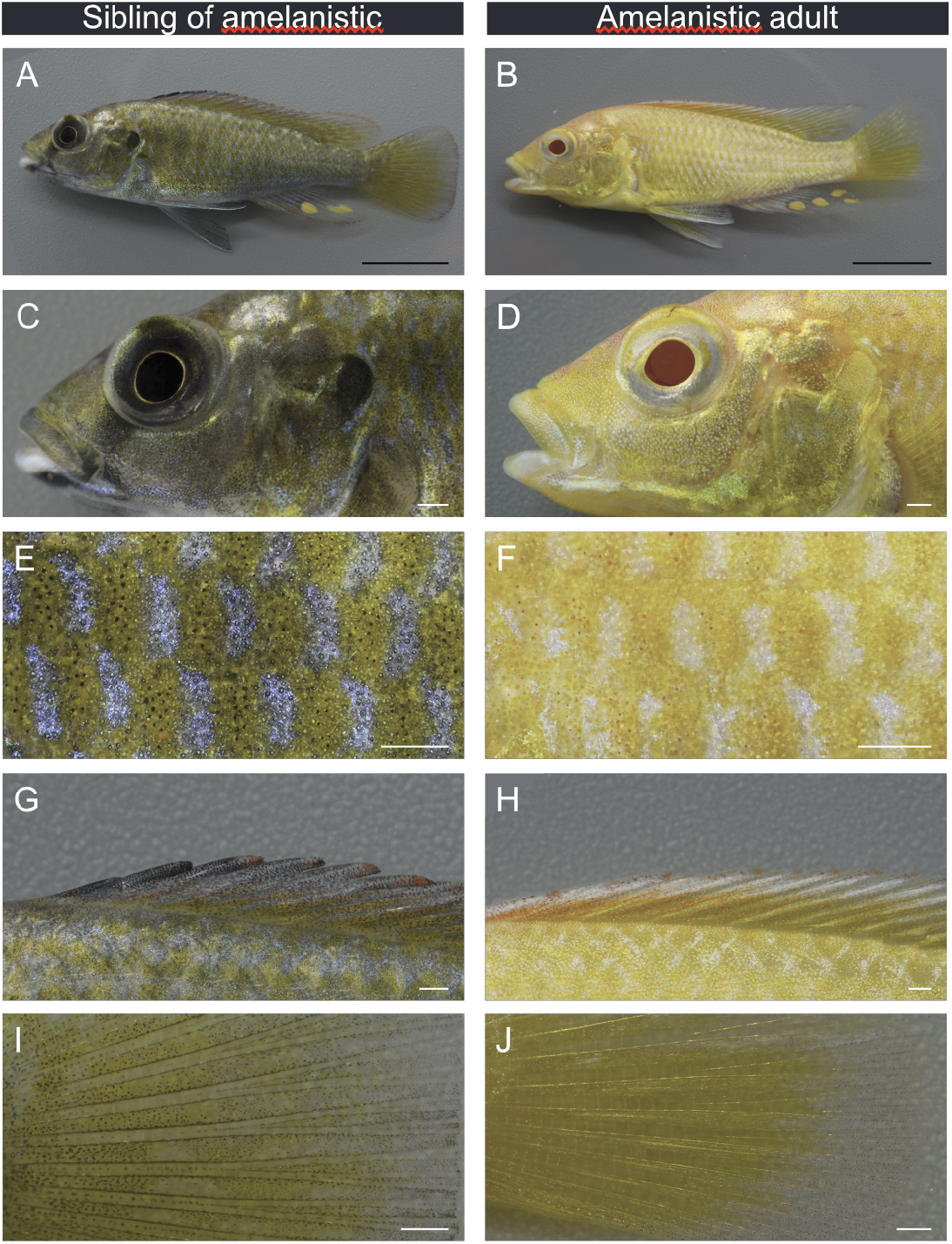
*oca2* amelanistic phenotype of adults. Comparison of F1s with wild-type and amelanistic appearance. **A-B)** full body **C-D)** head **E-F)** close-up of anterior trunk **G-H)** dorsal fin anterior **I-J)** caudal fin close-up of central spines, base of fin to the left and distal edge at the right. Scale bars 1 cm A-B and 1mm C-J.

## Discussion

Mapping genotypic variation to phenotypic variation is one of the major goals of evolutionary biology. Hence, a multitude of candidate loci underlying adaptive trait variation have been identified in a wide range of organisms (Courtier-Orgogozo et al., 2020). Most of these studies are performed in species harbouring natural variation but that are typically considered non-model systems due to the lack of tractable genetic tools. The recent development of genome editing tools such as CRISPR/Cas9 is thus revolutionising the field of evolutionary biology, allowing for candidate gene function tests in virtually any organism and uncovering the genomic and developmental basis of adaptation and diversification. In this study, we adapted existing protocols to establish CRISPR/Cas9 genome editing in the cichlid fish *Astatotilapia calliptera*. We generated both coding and non-coding mutations in *oca2* that were efficiently transmitted through the germline to the next generation. To our knowledge, this is the first report of successfully targeting gene function in this species or any cichlid within the Malawi radiation, and as such, represents the first step towards testing the genes and regulatory elements underlying variation in the Malawi cichlid radiation.

The amelanistic phenotype of *oca2* is expected given the role of *oca2* in tyrosinase transport for melanin production and melanosome maturation in melanophores (Beirl et al., 2014; Costin et al., 2003; Manga et al., 2001). This demonstrates that *oca2* is a useful first gene to target for establishing and refining CRISPR/Cas9 editing in a new species: it is easy to screen for mutations as with other melanin synthesis pathway genes (Li et al., 2021). Similarly to *Tyrosinase* knock-outs in the Lake Tanganyika cichlid *Astatotilapia burtoni* (Li et al., 2021), this *oca2* mutant line would permit unobstructed imaging of subdermal structures and fluorophores *in situ* during embryo development, enabling inter-specific comparisons for such studies.

Despite the lack of melanin, typical colour patterns are still noticeable on amelanistic *A. calliptera*. As teleost fish colour patterns self-organise through interaction between different pigment cells, this suggests that unpigmented melanophores are still present and contributing to pattern formation (Patterson and Parichy, 2019). However, *oca2*-deficient zebrafish also show an increase in iridophore numbers suggesting that the loss of *oca2* could affect other pigment cells (Beirl et al., 2014). We consider both of these possibilities when interpreting the amelanistic colouration phenotype. For example, the switch from blue to white reflective colouration in the alternating patches on the trunk indicates that black pigment is a component of blue colouration in *A. calliptera* (Figure 6). The mechanism may be similar to colouration in Siamese fighting fish (*Betta splendens*) where melanophores enhance the chroma and purity of the blue colour when underlying iridophores (Amiri and Shaheen, 2012). Alternatively or additively, the colour switch could be due to the loss of an influence of melanophores on iridophores, as zebrafish melanophores may induce iridophores to change shape and colour from white and dense to blue and loose (Owen et al., 2020). Similarly, the greater visible yellow/orange colouration due to xanthophores, red erythrophores, and bright reflective patches of iridophores in amelanistics could be due to melanin obscuring this pigmentation in wild-types (Figure 6). In some cichlid species superficial melanophores are found in the dermis, above the hypodermis (Beeching et al., 2013) so it is possible that they cover other pigment cells in *A. calliptera*. Alternatively, there may be greater numbers of these cell types in the amelanistic individuals. Comparison of the number and type of pigment cells during wild-type and amelanistic development may provide initial insights into chromatophore interactions in cichlids.

Here, we targeted coding and non-coding *oca2* sequences demonstrating that despite low embryonic survival, mosaic mutants and germline transmission occurs at an efficient rate. Embryo survival was low, averaging 20%, with most deaths occurring due to perforation of the yolk and its subsequent leakage. This low survival is comparable to the microinjection survival rates observed in other cichlids (20% in *Oreochromis niloticus* and 30% in *Astatotilapia burtoni*) (Li et al., 2021, 2014). Despite the low survival, mosaic mutant generation occurred at a high rate. Mosaic frequency was higher for coding sequence mutants (~50%) than for the non-coding deletion mutants (~12.5%). This likely reflects the lower efficiency of the HDR mechanism compared with NHEJ (Mao et al., 2008). Alternatively, this result may also reflect locus-dependent differences in mutation rate. To distinguish between the two hypotheses, a comparison between HDR and NHEJ modifications at the same locus is required. Nonetheless, we observed transmission of mutations to the next generation in both cases. *A. calliptera* species reach maturity on average at 8 months at which point they usually lay on average ~20 eggs, with clutch size increasing with age and size (Parsons et al., 2017). Despite low survival and low clutch sizes at young ages, germline transmission is high and as such it is possible to establish a breeding population of stable mutant *A. calliptera* within 16 months. One possibility to increase spawning frequency and increase clutch sizes to maximise the number of mutants, is peritoneal injections of Ovaprim, a commercially available mixture of gonadotropin-releasing hormone analogue and a dopamine receptor antagonist. Such injections resulted in a reduction in spawning interval by five days and a 2-fold increase in egg yield in *Astatotilapia burtoni* (Li et al., 2021). A similar effect would be expected in *A. calliptera.*

We were able to verify successful deletion of a ~1100bp stretch of the *oca2* 3′ UTR via HDR. However, whilst we could detect precise deletions in the G0 mutant and in two F1 individuals, some F1 progeny also contained a ~100bp indel. This insertion was likely the result of erroneous integration of fragments of the HDR template, as some regions shared homology with the template but in random orientations. Erroneous integration of HDR template fragments has been reported in CRISPR/Cas9-mediated HDR in other species including zebrafish, in which frequency of template integration was found to influence overall knock-in efficiency (Boel et al., 2018). In future applications, refinement of HDR template composition and chemical impairment of the NHEJ pathway may improve HDR efficiency in *A. calliptera* (Maruyama et al., 2015).

Genome editing in this species is particularly relevant on many fronts. First, this species is highly diverse, inhabiting a range of habitats (lacustrine and riverine) and showing extensive populational variation in several morphological, physiological and life history traits (Parsons et al., 2017). *A. calliptera* ‘masoko’ in particular, is a key example of ongoing sympatric speciation, with two divergent ecomorphs differing in depth habitat and dietary preferences and many other morphological traits, such as male colour, craniofacial profile and pharyngeal jaws. These differences are associated with assortative mating and local adaptation providing a good setup to address the early stages of adaptive diversification within the context of both natural and sexual selection. Further, there are plenty of genomic resources for this species, with a reference genome assembly at the chromosomal level and with hundreds of low coverage genomes distributed across several populations (Malinsky et al., 2018, 2015; Munby et al., 2021). These genomic resources combined with the genome editing tools and *A. calliptera* amenability to the lab allows for the tackling of adaptive diversification from both the genomic and developmental point of view. Additionally, it has also been suggested that the Malawi cichlid radiation initially stemmed from a generalist *Astatotilapia-type* lineage. The ~ 850 Malawi cichlid species can be grouped into seven eco-morphological groups, resulting from three separate cichlid radiations that stemmed from a generalist *Astatotilapia*-type lineage (Figure 1A) (Joyce et al., 2011; Malinsky et al., 2018). The divergence started with the split of the pelagic genera *Rhamphochromis* and *Diplotaxodon*, followed by the shallow- and deep-water benthic species, as well as the utaka lineage (water column shoaling cichlids), and finally the split of mbuna (rock dwelling cichlids). The ancestor of these three radiations was, most likely, very similar to *A. calliptera*, in terms of ecology and phenotype (Malinsky et al., 2018). As such, *A. calliptera* is a useful model in which to develop functional tools to explore Malawi cichlid explosive diversification.

An important attribute of Malawi cichlids is the ease of establishing inter-specific crosses for genetic mapping of traits of interest. Using such an approach, several studies identified genes associated with inter-specific variation in craniofacial profiles, jaw attributes, colour patterns and sex determination systems. The increase in genomic resources and the low costs of whole genome sequencing are also leading to an increase in genome-wide association studies in wild populations giving unprecedented insights into intraspecific variation (Kautt et al., 2020; Munby et al., 2021). A commonality between all these studies is that often the causal variants are in non-coding regions, hence establishing methods to edit non-coding regions will facilitate the dissection of their functional role. Here, both non-coding and coding sequence editing protocols are suited for loss of function experiments. The next step is to establish the targeted introduction of specific mutations using a knock-in approach, whereby a genomic variant associated with variation across species can be transferred from one species to the other. This approach will provide the causative link between genotype and phenotype variation and provide a genetic and developmental mechanism as to how organismal variation emerges.

## Conclusion

In summary, we have demonstrated the successful targeting of coding and non-coding sequences in the cichlid *A. calliptera* using CRISPR/Cas9. As the extant species of the lineage ancestral to the lake Malawi cichlid radiation, and as a very diverse species complex itself, *A. calliptera* is an ideal species with which to test hypotheses regarding speciation, adaptation and trait diversification. The establishment of genome editing tools for such key non-model species promises to reveal novel genetic and developmental mechanisms by which organismal diversity emerges.

## Supplementary Figures

**Figure S1:**
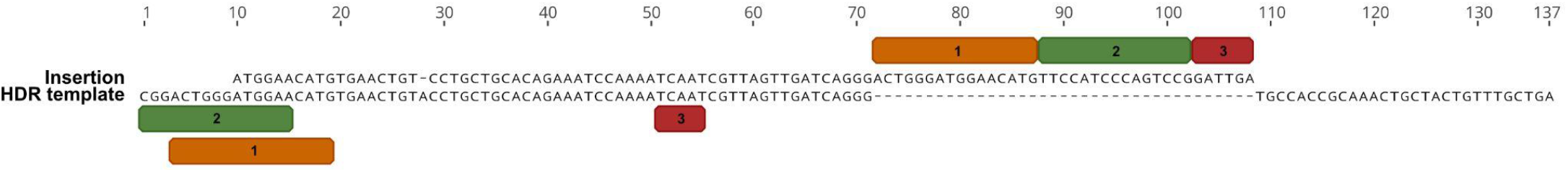
The HDR template and the insertion in *oca2* 3′ UTR mutants (F1_#5, F1_#7 and F1_#13) shows a high degree of homology. Part of the insertion that does not align with the HDR template sequence (coloured boxes 1, 2 and 3) shows homology with different upstream regions of the HDR template (coloured boxes 1, 2 and 3).

**Figure S2:**
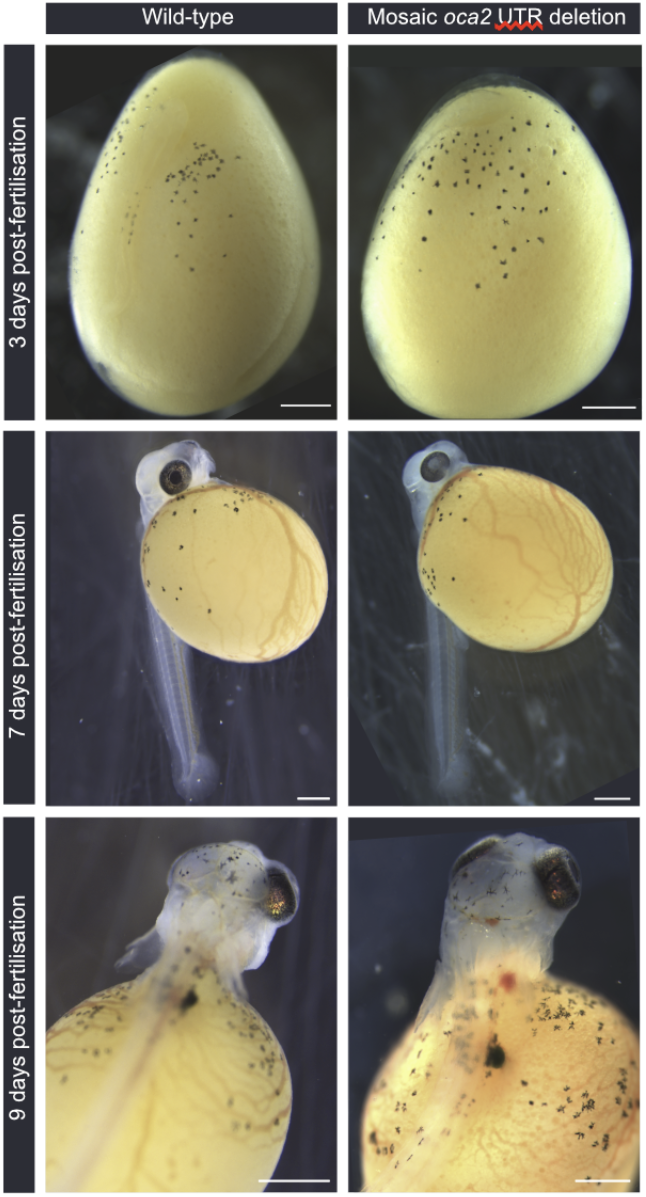
*oca2* 3′ UTR embryonic G0 phenotype.

**Figure S3:**
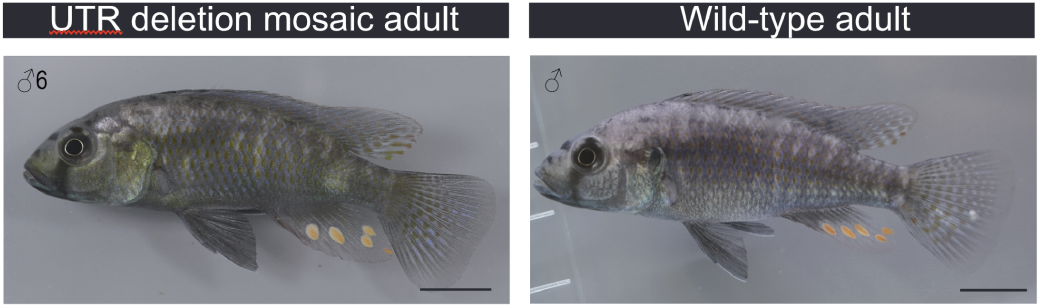
*oca2* 3′ UTR G0 adult phenotype. Numbering corresponds to table 4.

## Methods

### Fish maintenance and crossing

*Astatotilapia calliptera* were kept under constant conditions (28 ± 1°C, 12 h dark/light cycle, pH 8) in 220 L tanks. All animals were handled in strict accordance with the protocols listed in the Home Office project licence PCA5E9695. Fish were fed twice a day with cichlid flakes and pellets (Vitalis). Tank environment was enriched with plastic plants, hiding tubes, and tank bottoms were covered with sand. Males were provided with a clay pot in which they established a territory and spawned with gravid females. Males and females were housed in the same tank but separated by a divider to control the timing of spawning. Males were housed singly, while females were kept in groups of 8-15 females. On the day of spawning, the divider was removed, and interactions were monitored for spawning. If spawning was detected, the fish were given an additional 30 - 60 minutes to fertilise the eggs. The fertilized eggs were then removed from the female’s buccal cavity and injected with sgRNAs and Cas9 protein.

### gRNA design and synthesis

CRISPR/Cas9 targets were selected with the CHOPCHOP software online (http://chopchop.cbu.uib.no/) using the *Astatotilapia burtoni* genome as a reference. Basic local alignment search tool (BLAST) (Altschul et al., 1990) was then performed with the *A. calliptera* genome at Ensembl to confirm homology and avoid off-targets. sgRNAs used in this study start with GG or GA followed by N18, which are directly upstream of the NGG PAM sequence (5′-GG-N18-NGG-3′ or 5′-GA-N18-NGG-3′) to satisfy the requirements for *in vitro* transcription using a T7 or SP6 promoter respectively. We designed three sgRNA in exon 1 (GACGGCATCCCAAGGCCACC, GGTCACCGAAGGCGGTGGCA and GGGGAACTATGTCTGCTGGA) one sgRNA in exon 3 (GAACAACGGCTCCCTGGACG) and two sgRNA in the UTR region (GAGTGGTCACACAGTTTCTT and GATCAACTAACGATTGATTA). The PCR primers for sgRNA synthesis are given in table S1. To synthesize sgRNAs we used the cloning free method described in Varshney et al. 2015 using T7 or SP6 Polymerases (NEB) depending on the 5′ sgRNA sequence. The sgRNAs were purified using the Qiagen RNeasy kit.

**Table S1:**
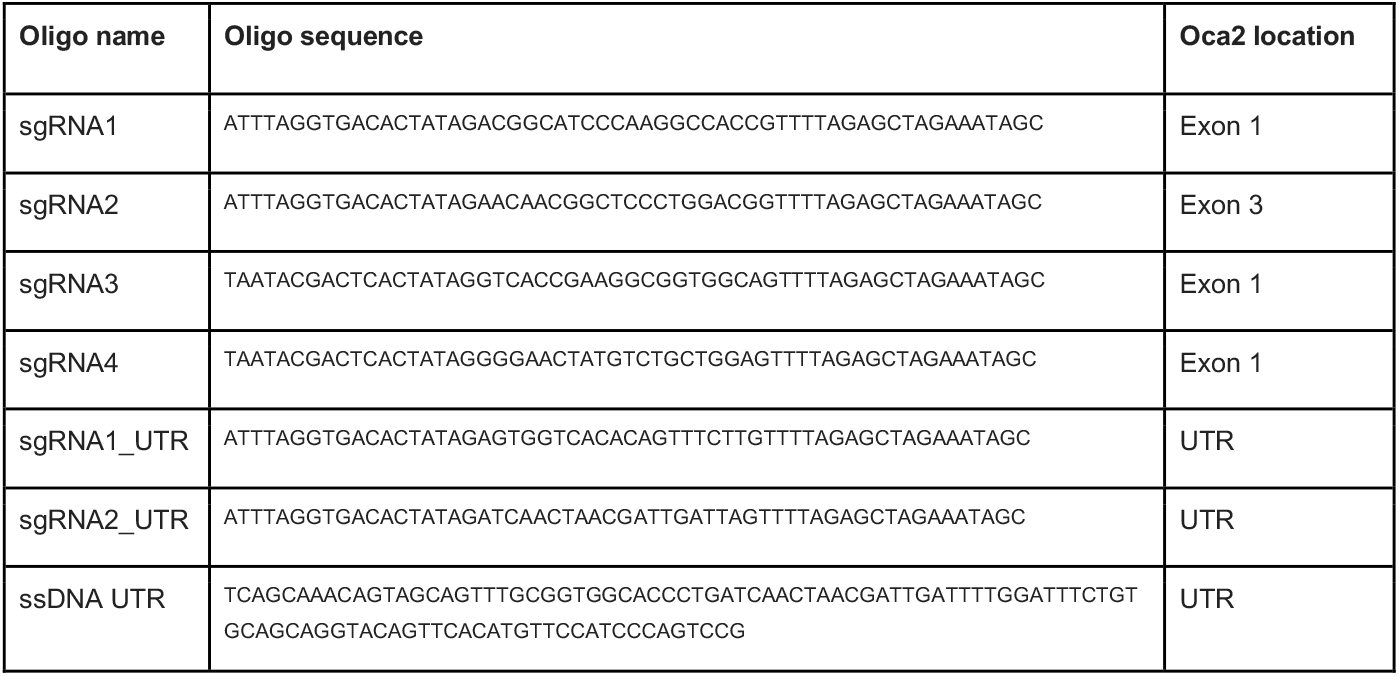
Oligo sequences used for sgRNA synthesis and ssDNA sequence.

### Microinjection

After collection eggs were placed into wells created by a mold with circular indentations in 2% agarose made with tank water see Li et al. 2021 for more information on the mold). Single cell embryos were injected with a mixture of sgRNAs at 300 ng/μl each, together with True Cut Cas9 Protein V2 (Invitrogen) at 150 ng/μl and dextran labelled with TexasRed (ThermoFisher Scientific, 10,000 MW) at 0.25%. Three injection mixes were used: 1) sgRNA1 and sgRNA2 targeting exons 1 and 3; 2) sgRNA3 and sgRNA4 targeting exon 1; or 3) sgRNAUTR1 + sgRNAUTR2. To improve deletion efficiency, a 100bp ssDNA (IDT Technologies) with left and right homology arms (Figure 3A) located at the outer sides of the Cas9 cutting sites was used at 20 ng/ul. Microinjection needles were pulled manually from glass capillaries (GC100F-10, 1.0mm O.D; 0.58 mm I.D, Harvard Apparatus) using a Sutter P-97. Needles were opened by gently tapping the needle on a Kimwipe to break the tip to a diameter of ~10 μm diameter. Each egg was injected using a pulse-flow nitrogen injection system (MPPI-3 with a back pressure unit) with 2 pulses at 1 ms and 40 psi (~1-2 nl). The injected embryos were kept individually in 6 well plates, in an orbital shaker at 25°C in the presence of methylene blue (10 mg/ml) and with daily water changes.

### Germline transmission rates and F1 progeny genotyping of *oca2* coding mutants

Four mosaic oca2 mutant males were reared until adulthood and crossed with wildtype females (Table 2). Germline transmission rates were quantified by genotyping potential F1 heterozygotes. DNA was extracted from 6-14 dpf embryos (after yolk removal) using the DNA miniprep kit (Zymo). PCR products were amplified with Phusion (NEB), following the manufacturer’s specifications, with an annealing temperature of 62 °C. Primer sequences for exon 1 and exon 3 genotyping are listed in Table S2. PCR products were purified with QIAquick PCR Purification Kit (Qiagen). Presence of heterozygous mutants was then confirmed using Sanger sequencing. Sequence analysis was performed using the Synthego ICE CRISPR analysis tool (https://ice.synthego.com/). This tool infers CRISPR edit sites from sequences derived from heterozygous or mosaic individuals. A summary of the analysis of Sanger sequencing fragments is detailed in Supplementary File 1. Sequence traces were analysed on Geneious Prime to detect sequence quality drops associated with the sgRNA cut site. Further, mutant sequences were extracted using the ICE CRISPR analysis tool by selecting the most frequent mutant allele and aligned with the MAFFT alignment plugin on Geneious Prime (Figure 2B, C and D). Two *oca2* mosaic coding mutant females were incrossed with one *oca2* mosaic male to generate F1s with two oca2 mutant alleles. Germline transmission was inferred by visual quantification of the number of embryos lacking melanic pigmentation. These estimates probably represent an underestimation of transmission rates, since heterozygotes do not present an amelanistic phenotype.

**Table S2:**
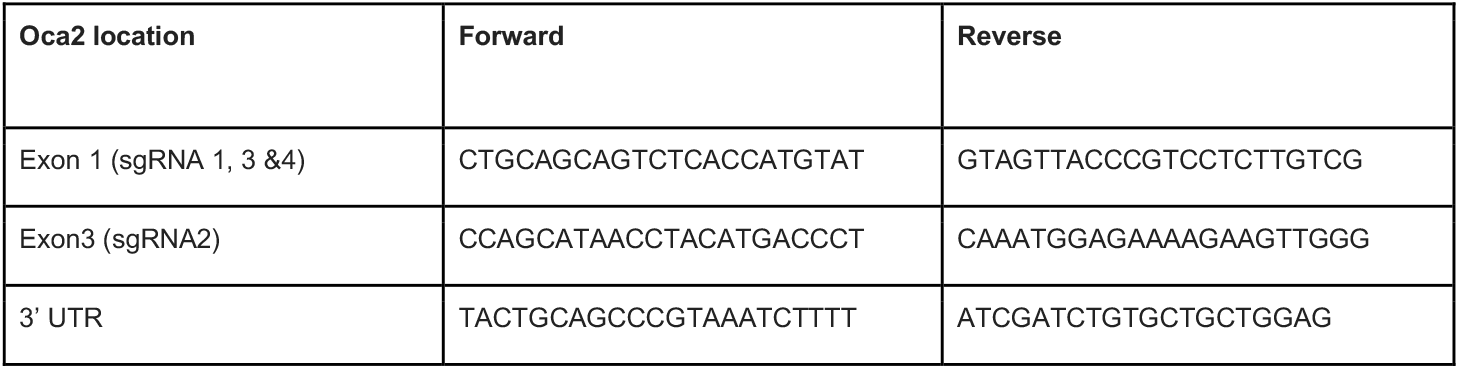
PCR primers for genotyping.

### Germline transmission rates and F1 progeny genotyping of *oca2* 3′ UTR deletion mutants

G0 mosaics and F1 heterozygous mutant progeny were assessed by PCR using two primers flanking the cut sites. The forward primer is located 207 bp upstream the left cut site and the reverse is 400 bp downstream of the right cut site (Table S2). This assay differentiates between the wild-type individuals and individuals carrying the desired deletion. PCR on wild-type individuals results in only one PCR fragment, whereas mosaic individuals carrying the deletion will show two fragments - the wild-type sequence (~1715bp) and the sequence containing the deletion (~624 bp). Only one G0 individual tested positive for the deletion. This G0 individual was crossed with wildtype females (Table 4). DNA was extracted from a fin clip of the G0 founder and from 8dpf F1 embryos (after yolk removal). PCR was performed with OneTaq (NEB), following the manufacturer’s specifications, with an annealing temperature of 60 °C. PCR purification of the deletion fragments was performed with QIAquick Gel Extraction Kit (Qiagen). Presence of the deletion in the G0 and F1 individuals was then confirmed using Sanger sequencing. Sequences were aligned using the MAFFT alignment plugin on Geneious Prime (Figure 3B).

### Embryo and adult imaging

Embryos were imaged on a Leica M205 FCA stereoscope with a DFC7000T camera under reflected light darkfield. For each embryo, images were taken at multiple focal distances. These images were then focus-stacked using Helicon Focus or Photoshop to produce a single image with all cells in focus. To prevent movement between imaging different focal planes, post-hatching embryos were anaesthetised by submersion in 0.02% Tricaine methanesulphonate (Sigma-Aldrich E10521) for the duration of imaging (approximately 2 minutes per embryo) with the yolk supported in a shallow well of solidified 1% low-melting agarose (Promega, V2111). Adult fish were photographed using a Panasonic DMC GX7 camera with a Panasonic Lumix G 20mm pancake lens, in a photography tank containing a scale. Lighting conditions were standardised using two light sources, one either side of the camera, and a grey background.

## Author contribution

BC, JE, AM and MES performed experiments and analysed the data. GFT, AMS and DJ contributed with fish stocks and initial experiment setup. MES, EAM and SAJ designed, provided resources and supervised the study. BC, JE and MES wrote the manuscript with contributions or feedback from all authors. All authors read and approved the final version of the manuscript.

## Additional information

Supplementary File 1 contains information on the analysis of the mutant sequences.

## Acknowledgements

We thank the Animal Technicians and NACWO at the UBS Cichlid fish facility in Madingley (Cambridge) for the help in rearing wildtype and mutant fish. Members of the Miska lab and Morphological Evolution Lab for technical support and discussion. BC and AM are supported by the Wellcome Trust PhD Programme in Developmental Mechanisms (222279/Z/20/Z and 102175/Z/13/Z respectively). E.A.M. is supported by Cancer Research UK (C13474/A18583, C6946/A14492) and the Wellcome Trust (219475/Z/19/Z, 092096/Z/10/Z). SAJ is supported by the HFSP grant RGY0079/2018 and the NSF grant IOS-1825723. ES is supported by a NERC IRF NE/R01504X/1.

